# Optimal Phylogenetic Reconstruction of Insertion and Deletion Events

**DOI:** 10.1101/2024.01.24.577130

**Authors:** Sanjana Tule, Gabriel Foley, Chongting Zhao, Michael Forbes, Mikael Bodén

**Affiliations:** School of Chemistry and Molecular Biosciences, The University of Queensland, Brisbane, QLD 4072, Australia; School of Mathematics and Physics, The University of Queensland, Brisbane, QLD 4072, Australia

## Abstract

Insertions and deletions (indels) influence the genetic code in fundamentally distinct ways from substitutions, significantly impacting gene product structure and function. Despite their influence, the evolutionary history of indels is often neglected in phylogenetic tree inference and ancestral sequence reconstruction, hindering efforts to comprehend biological diversity determinants and engineer variants for medical and industrial applications.

We frame determining the optimal history of indel events as a single Mixed-Integer Programming (MIP) problem, across all nodes in a phylogenetic tree adhering to topological constraints, and all sites implied by a given set of aligned, extant sequences. By disentangling the impact on ancestral sequences at each branch point, this approach identifies the minimal indel events that jointly explain the diversity in sequences mapped to the tips of that tree. MIP can recover alternate optimal indel histories, if available.

We evaluated MIP for indel inference on a dataset comprising 15 real phylogenetic trees associated with protein families ranging from 165 to 2000 extant sequences, and on 60 synthetic trees at comparable scales of data and reflecting realistic rates of mutation. Across relevant metrics, MIP outperformed alternative parsimony-based approaches and reported the fewest indel events, on par or below their occurrence in synthetic datasets. MIP offers a rational justification for indel patterns in extant sequences; importantly, it uniquely identifies global optima on complex protein data sets without making unrealistic assumptions of independence or evolutionary underpinnings, promising a deeper understanding of molecular evolution and aiding novel protein design.

## 1 Introduction

Insertions and deletions (indels) play a pivotal role in shaping the landscape of genetic diversity. The importance of indels in protein families lies in several key aspects: 1) indels can introduce variation in the length and composition of protein sequences, leading to anything from subtle to dramatic changes in protein structure and function, which in turn arm us to engineer novel proteins suited to medical and industrial applications [17, 24, 25, 33, 12]; 2) indels serve as key evolutionary markers, offering insights into the history of and relationships among homologous sequences [19, 30, 4]; 3) indels probe and inform the relationship between the multiple sequence alignment (MSA) and phylogenetic tree inference, since indels are much less homoplastic than substitutions [15, 5]. To ensure sequence characters line-up to indicate homology, the MSA uses gap characters to pad where deletions and insertions could have occurred in the evolutionary past, forming patterns that can only be resolved in relation to other included sequences.

A key difference between methods that account for or infer indels is how they represent their occurrence in extant sequences of a family, and then how they are encoded at ancestral branch points. That said, it is not unusual to simply treat indels as missing data [34, 13, 29, 21].

A *position-specific* (PS) discrete encoding (often binary) assumes that (like any other character in an alignment) indels occupy a single site and does not allow indels to overlap or be subsets of other indels [6, 18]; this implies that calls of one site are independent of other sites in a molecular sequence, so finding the position-specific optimum is part of the global optimum [31, 32, 1, 23, 3]. The problem with defining the global optimum this way is that both insertions and deletions can involve more than one consecutive position.

A *composite* discrete encoding, such as simple indel coding (SIC) [26], represents *an indexed set of* indels that occur with different lengths and at different, potentially overlapping positions; with SIC, for any sequence across the input data, an indel is either present, absent, or inadmissible, which is used to flag when it is impossible to say, e.g. a deletion is precluded by an earlier, wider deletion [26]. A discrete state at each index (or position) lends itself in general to a multitude of inference algorithms, including maximum-likelihood (ML; provided a probabilistic model of transitions between the states) and maximum-parsimony. Indeed, FastML [22] offers both for SIC, utilising a gene loss/gain model for ML.

Partial order graphs (POGs) can represent a family of sequences via nodes for the *inclusion* of homologous content (using a global alignment index) and directed edges to selectively indicate the *exclusion* of content in other sequences [9]. A single sequence can be generated by following edges from a global “dummy” start to a “dummy” end point; we consider each sequence as having an indel “footprint” as represented by the set of edges that are followed. As in the ancestral sequence reconstruction method GRASP [7], we formulate the indel inference problem over the set of all edges. Each global alignment index is associated with a finite and discrete set of edges (each of which has a start and end index in turn), enabling the use of both ML and parsimony-based inference to chose which edge to traverse at each index. In contrast to composite discrete encodings, edges can capture some positional dependencies, given that edges are chosen to the exclusivity of others that share one of the two alignment indices. Most existing indel inference methods solve a fragmented optimisation problem, to be resolved at each internal node rather than a global problem across the phylogenetic tree. GRASP [7] uses a method referred to as bi-directional edge parsimony (BEP; or alternatively maximum-likelihood; BEML) that finds “optimal” edges, but not optimal paths through them, failing to provide guarantees of the optimal composition of indels within each (ancestral) sequence; a solution is therefore not guaranteed to be globally optimal.

When indel dependencies and whole-sequence constraints are considered, the indel inference problem becomes non-trivial [8], and stymie probabilistic adaptations. Hence, we focus on parsimony-based formulations that do not account for evolutionary distances.

Fredslund and colleagues [8] address the presence of long indels (exceeding one site) by using the concept of a gap graph with an affine gap penalty function. Snir and Pachter [27] employ a dynamic programming approach to deduce the insertion and deletion history in long DNA sequences within the context of small species phylogenetic trees. The Snir and Pachter (SP) recursive dynamic programming method determines whether there is a gap or non-gap at the current position in the molecular sequence at a node. Reassuringly, it takes into account the previous position and the parent of the node being considered, but it comes with a time complexity exponential to the number of nodes in the tree, and this complexity is likely to hamper applications to protein datasets.

In this paper, we hypothesise that framing the indel inference problem as a single global optimisation problem across all branch points in a phylogenetic tree accounting for multi-site dependencies amongst indels can lead to qualitatively and quantitatively better reconstruction of their evolutionary history relative to existing parsimony-based approaches.

We model indel inference in the phylogenetic tree as a maximum-parsimony problem using edges to explicitly represent multi-site dependencies amongst indels, as imprinted on the input sequence alignment when mapped to the phylogenetic tree. Our methodology draws inspiration from MSA algorithms, utilizing gap opening and pattern differences between two sequences as the global optimisation objective. The constraints and objective function can be readily represented as a Mixed-Integer Programming (MIP) problem. The resulting MIP has a large embedded network flow component together with side constraints used to evaluate the objective. Our working hypothesis was that this MIP is tractable to solve, even if it had millions of variables and constraints.

Our assessment of indel solutions involves both quantitative and qualitative analyses, comparing them with other parsimony-based approaches based on three criteria: 1) total indel events, 2) evolutionary cohesiveness, and 3) indel footprints. For synthetic data, we are also able to compare the number of predicted indels to the actual number of indels introduced during evolution.

## 2 Methods

### Preliminaries

A phylogenetic tree *T* = (*X, N, B*) is a directed acyclic graph (DAG) that represents the evolutionary relationships *B* among a set of leaf (or extant) nodes *X* (indexing extant sequences in the alignment) and internal nodes *N* (indexing the ancestors of *X*). Assuming we have *n* extant nodes then, as the tree is binary, we have *n* − 1 internal nodes. By convention we denote *N* = {1, …, *n* − 1}, with 1 as the root node, and *X* = {*n*, …, 2*n* − 1}. For each branch (*k, l*) ∈ *B* (where *k* is the direct ancestor of *l*) we have *k* ∈ *N, l* ∈ *X* ∪ *N* and *k* < *l*. A multiple sequence alignment of extant nodes *X* is a mathematical representation of the arrangement of homologous molecular sequences from different species or sources. If the sequences are of length *m* we denote the sites in the sequences as *S* = {1, …, *m*}. We encode the MSA using a matrix of binary data *A* = *a*_*ki*_, *k* ∈ *X, i* ∈ *S* where 0 represents a gap and 1 any amino acid at the relevant site.

We define *P*, a partial ordered alignment graph (POAG), as the union of the edges between adjacent occupied sites in *M*. More formally *P* = ∪_*k*∈*X*_{(*i, j*)|*i, j* ∈ *S, i* < *j, a*_*ki*_ = 1, *a*_*kj*_ = 1, *a*_*ks*_ = 0 *∀ i* < *s* < *j*}. We denote the set of edges in *P* that arrive at site *i* as 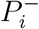 and the set of edges that leave site *i* as 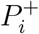. *P* captures the relationship between non-gapped positions across all sequences in the MSA, and we refer to this as the indel footprints in a given set of extant nodes *X*. Figure 1 shows a small MSA and its conversion to a POAG.

**Figure 1:**
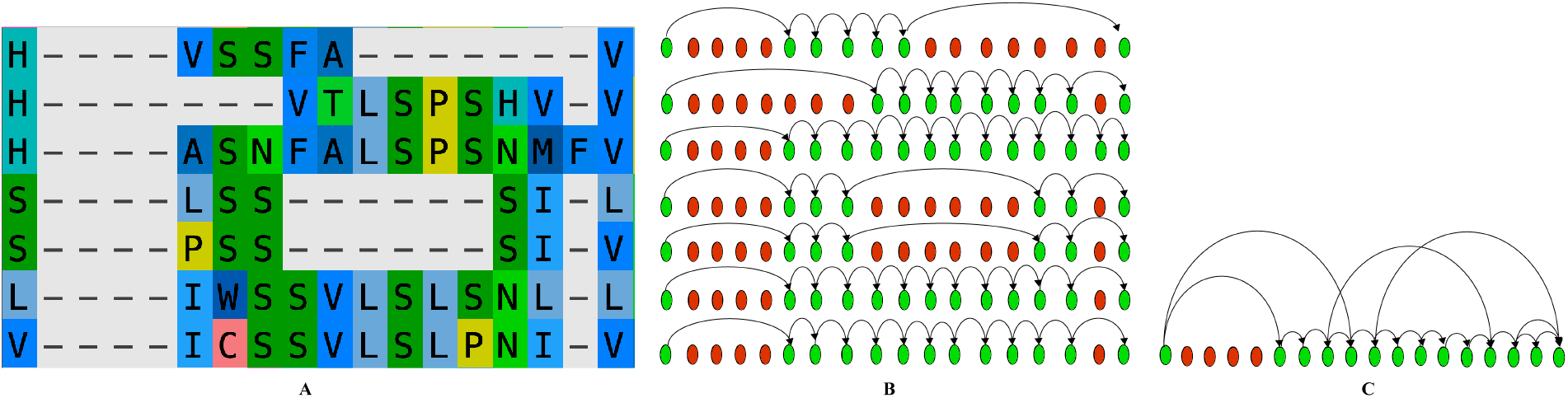
A) An example MSA; B) POGs from the example MSA, where green and red nodes represent non-gapped and gapped positions in a sequence, respectively; C) Overall POAG representing the union of edges present in POGs.

The first and last site in each sequence can be gapped, so we introduce “dummy” start and end nodes in the POAG. We can, without loss of generality, preprocess our data to remove any consecutive sites that are gapped and any consecutive sites that are non-gapped in every sequence.

We focus on optimising the indel events for a phylogenetic tree and an MSA using a MIP formulation where *T, A* and *P* are the input data structures. We compute the optimal path of edges forming a sequence for each of the nodes in *N* while restricting the indels used to those implied by the edges in *P*. For computing the optimal path, we use variables *d* as difference in position wise indel pattern and *g* as gap opening between two indel patterns.

The MIP uses the following decision variables:

- *a*_*ki*_ ∈ {0, 1} encodes the gap or non-gap position at internal node *k* ∈ *N* at site *i* ∈ *S*. Note that *a*_*ki*_ is input data for *k* ∈ *X*, but a variable for *k* ∈ *N*.
- *y*_*kp*_ ∈ {0, 1} encodes the presence or absence of edge *p* ∈ *P* for internal node *k* ∈ *N*.
- *d*_*bi*_ ∈ {0, 1} is 1 if the indel pattern is different for the two nodes in branch *b* ∈ *B* at site *i* ∈ *S*. These variables are used to count site differences between neighbours in *T*.
- *g*_*bi*_ ∈ {0, 1} i.e. gap opening is 1 if a difference starts for the two nodes in branch *b* ∈ *B* at site *i* ∈ *S*. These variables are used to count the number of sequences of differences between neighbours in *T*.

We define MIP-Indel as:
min

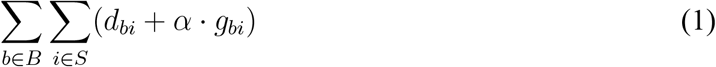

s.t.

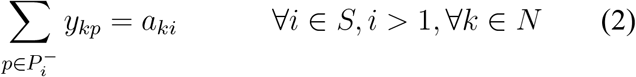

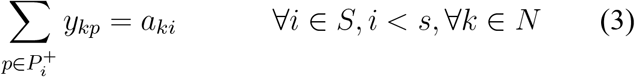

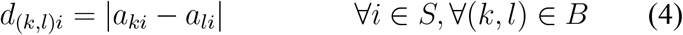

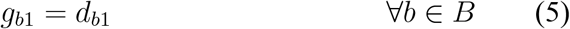

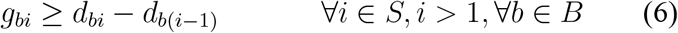

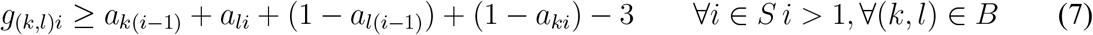

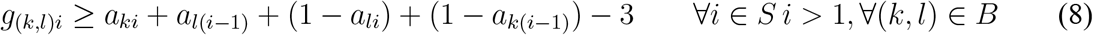

The objective value (1) minimises a weighted combination of the number of individual site difference and the number of sequences of site differences. Constraints (2) and (3) ensures that site gap/non-gap indicator is equal to the total edges leaving and arriving at the site. Constraint (4) computes the site difference between the two ends of a branch in the tree. Constraints (5-8) compute the number of times sequence differences occur. Constraint (6) detects the beginning of series of differences - the sequences have been the same, but now they are different. Constraint (7) and (8) count the case where the sequences are different and stay different, but the gap opening patterns reverse. *α* ∈ ℝ^+^ is the hyperparameter representing the importance of minimizing opening a new gap between indel patterns in two molecular sequences. The implementation is available via GitHub at https://github.com/santule/indelmip.

### Datasets

We determine the characteristics of predictions and computational performance of methods for indel inference using both real (but imperfectly aligned) protein families as well as sequence data that was generated by simulating protein evolution (resulting in perfect alignment). The latter provide us with opportunity to evaluate the ability of methods to reconstruct the synthetic history of mutations and determine the robustness at a greater level of granularity and control than is practical with biological data.

We use 15 real datasets from a varied range of protein families, with total extant sequences ranging from 165 to 2000 and aligned sequence length of protein sequences ranging between 513 and 5240. The real datasets are denoted by combining a brief protein family name with the total number of existing sequences. For instance, the dataset RNaseZ_624 comprises sequences of Metallo-*β*-lactamase family, encompassing a total of 624 protein sequences. For details on the datasets, please refer supplementary section 1.1.

For synthetic datasets, we generate trees for *n* ∈ {300, 500, 700, 1000, 2000} extant sequences with a mean evolutionary distance *δ* ∈ {0.02, 0.5, 0.8, 1} from a single root using the tool TrAVIS, which forms part of the GRASP-suite [7]. TrAVIS samples the Gamma distribution *γ*(*κ, θ*) (where *κ* ∈ {0.5, 1, 2} defines the shape and *θ* = 0.2 the scale) to set distances on each branch (in turn normalised to the mean *δ*), bi-furcating each branch point until the specified number of sequences have been mapped as leaves. Details on synthetic data generation using TrAVIS is covered in supplementary section 1.2. Synthetic datasets are named by concatenating *n, δ* and *κ*. For example, `t300d0.02s1` means a tree with 300 leaves with a mean distance of 0.02 to the root, generated using a Gamma shape parameter of 1.

## 3 Results

We compare our MIP Indel method with other methods for indel inference, including BEP, PSP (parsimony method based on PS indel encoding) and SICP (parsimony method based on SIC indel encoding) [22]. We also compare against Snir and Pachter’s method that (similar to our approach) uniquely attempts to identify a globally optimal parsimonious solution.

The MIP model is solved using Gurobi [10], the BEP, SICP, PSP are run using GRASP [7]. For MIP, the value of hyperparameter *α* (to weight gap introduction and extension) is set to 2 (see Supplementary Section 1.5 for an exploration of *α* settings). For BEP, SICP and PSP, we use the sequence (path through a POG) that GRASP produces by default for each branch point composed from multiple optimal solutions. In turn, this implies that indel events between ancestor sequences can originate from different (equally optimal) solutions for the same input data.

All the experiments were run on Dell PowerEdge R630 2 × 2.6 GHz 14-core Xeon (28T) with 256 GB RAM, using a single thread with one hour total time limit per dataset. We attempted to make inferences on 15 real and 60 synthetic datasets; most methods returned solutions, but the SP method scaled poorly with tree size and was unable to complete analysis on any real data set.

### 3.1 MIP Indel infers least number of indel events

Post-inference, each branch (*k, l*) ∈ *B* (where *k* is the direct ancestor of *l*) implies a minimum number of indel events. Equation 9 breaks down the binary differences in (continuous) gap states between *k* and *l*.

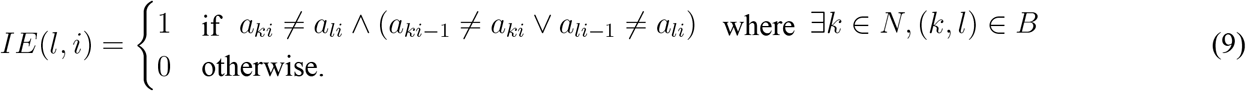

The indel score across the tree is defined by Equation 10.

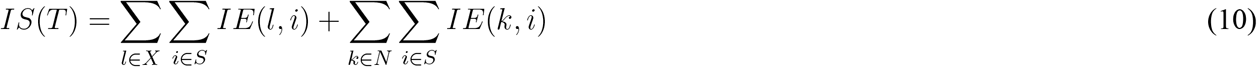

Figure 2 shows an example of how indel events are determined and how the indel score is calculated for a minimal example tree.

**Figure 2:**
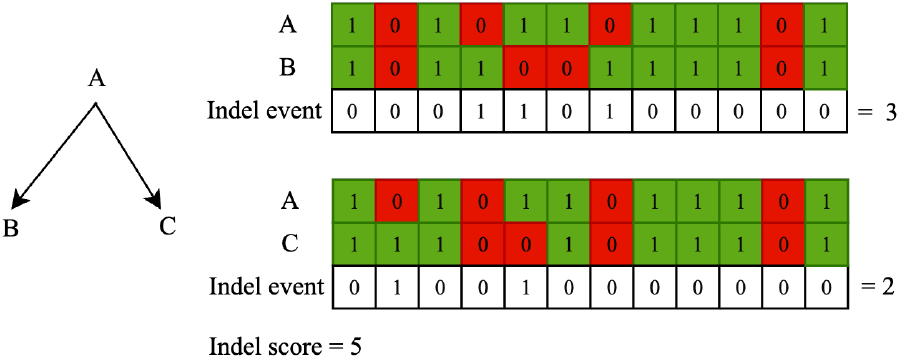
Example of indel events and score for an example tree. A is ancestor of B and C. 1 refers to non-gap and 0 refers to gap in molecular sequence respectively.

The indel score for the 15 real datasets are shown in Figure 3A. In all datasets, MIP Indel reports least indel score, followed by PSP, BEP and SICP methods. The RNaseZ_624 dataset represents a large, ancient protein family and commensurate sequence divergence is particularly challenging; for this data, all methods reported the greatest number of indels, and the BEP method failed to return any results within the time limit. The DHAD_1658 dataset is slightly less complex, and recapitulated the overall trend that MIP finds a solution with the smallest number of insertions and deletions, ahead of baseline position-specific (maximum) parsimony. Results for synthetic datasets in general corroborate the trends for the real data (see Supplementary Figure 1.4).

**Figure 3:**
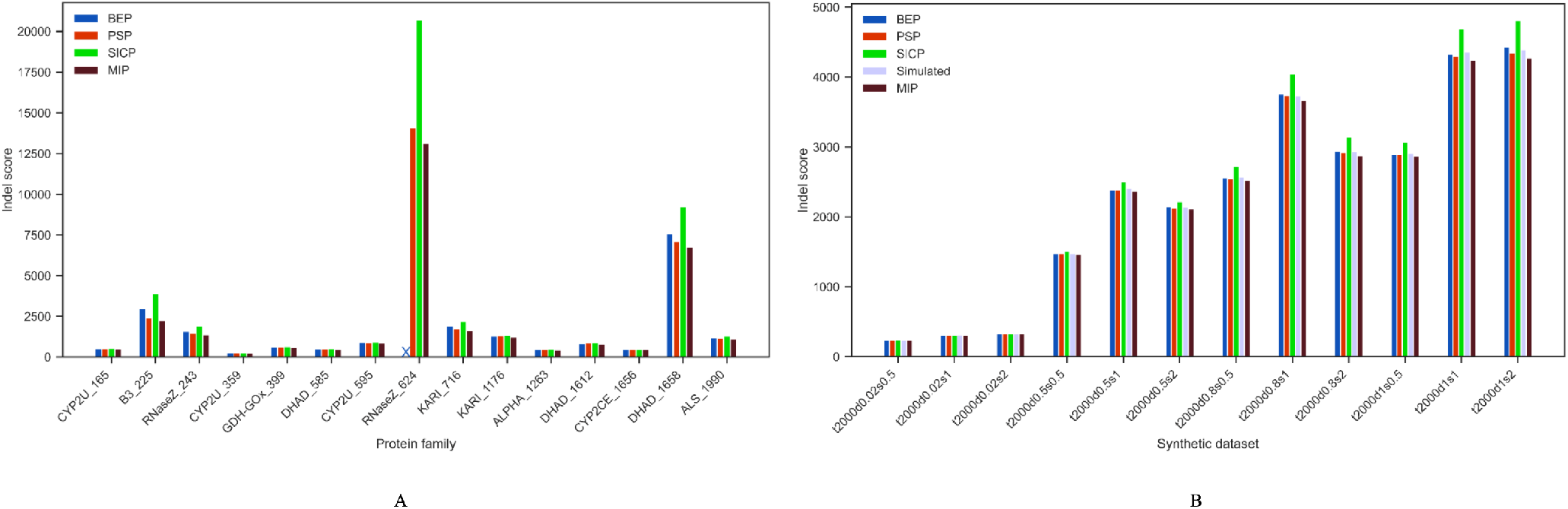
A) Indel score for 15 real datasets. X in the above figure denotes that BEP did not provide a solution for the dataset RNaseZ_624; B) Indel score comparison between (synthetic) ground truth and MIP Indel solution for trees with 2000 extant sequences with varying mean distance *δ* and *κ*.

We also compared MIP with the SP method on the smallest synthetic datasets (see Supplementary Section 1.6 on how these small synthetic datasets are generated). The SP method is unable to finish analysis on any of the other realistically large datasets; when completed, SP identified an equal or slightly greater number of indel events (see Supplementary Figure 6).

For the 60 synthetic datasets, we have access to the intermediate, ancestral sequences. From these sequences, we derive indel patterns by assigning the values of 1 and 0 to gap and non-gap positions, respectively. Subsequently, employing the methodology outlined in Section 3.1, we calculate the “ground truth” indel score. Figure 3B presents a comparative visualisation of the ground truth indel score alongside the indel scores obtained through various methods for distinct trees featuring 2000 extant sequences.

In instances where the mean distance within the tree is minimal, the MIP indel score strongly correlates with the ground truth indel score. Conversely, as the mean distances become larger, the ground truth score exceeds the MIP indel score. This outcome is anticipated, as greater distances elevate the likelihood of greater number of indel events between branch points. On the other hand, the SICP method scores higher than the ground truth as the mean distances become larger. Both BEP and PSP solutions closely align with the ground truth, with BEP method occasionally scoring higher than the ground truth.

### 3.2 MIP Indel solutions are evolutionarily cohesive

In contrast to methods that do not determine global optimas, and methods that assume degrees of site independence, MIP offers the distinct advantage in enforcing decisions that rationalise the *global* history of events. Across a tree there are scenarios of insertions and deletions that should not occur at all, e.g. a deletion event at a position that was inserted in an ancestor and subsequently deleted in its descendants. Additionally, there are mutational events that are highly unlikely to occur, e.g. a deletion occurring at the identical position in branch points directly descending from a common ancestor where content was inserted from its own ancestor.

We define evolutionary cohesiveness by observing the changes in indel states *between* branch points. We quantify the number of branch points possessing at least one site where the predicted indel state differs completely from that of all its neighbors in the tree; specifically, a site being gapped at the branch point, when all its descendants and its ancestor are non-gapped, and vice versa. Illustrative scenarios are presented in Figure 4A, where 0 and 1 represent a gapped and non-gapped state at a sequence site, respectively. This measure of cohesiveness holds significance in ascertaining the level of overall heterogeneity of indel evolution in a phylogenetic tree, indicative of an interchanging history. We hypothesised that the level of cohesiveness of MIP indicates greater level of homogeneity relative to alternative methods. Indeed, MIP infers *no* such ancestors across all real and synthetic datasets (see Figure 4B), except for one site involving one ancestor in the case of RNaseZ_624.

**Figure 4:**
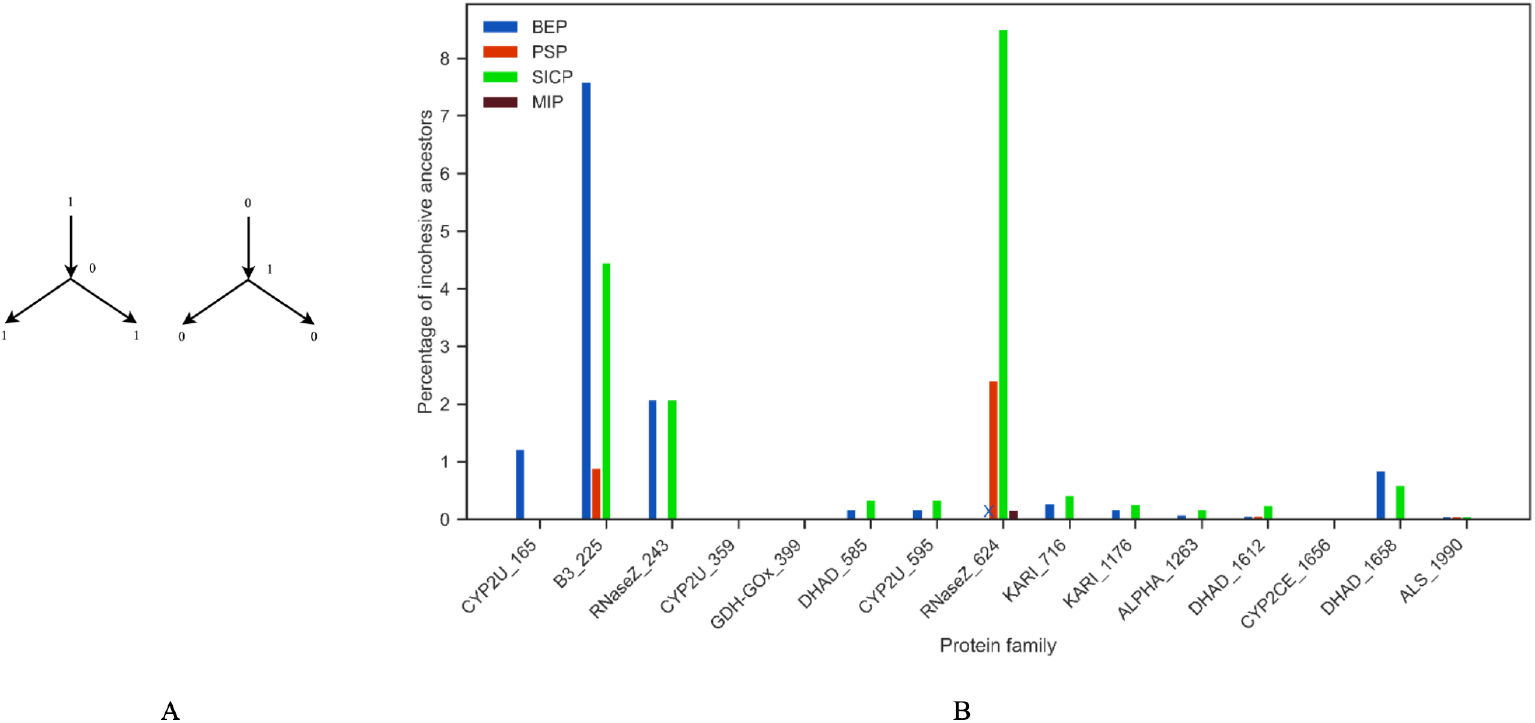
A) Example of “incohesive” indel states at a site at an ancestral branch point, where the state is distinct from both its descendants and its ancestor. B) Percentage of incohesive ancestors for 15 real datasets. The symbol ‘X’ in the figure indicates that BEP did not find solution for the RNaseZ_624 dataset.

Both BEP and SICP methods fall short in cohesiveness for certain datasets, which is unsurprising given that they optimise indel events locally and/or do not account for dependencies. PSP appears to introduce a small number of incohesive branch points, but we explain this by the occurrence of multiple (equally optimal) solutions and that the implementation we use mixes how they are used across different ancestors. We expect that if we stopped at a single optimal solution, the column-specific cohesiveness for BEP and SICP would be improved slightly and be perfect in the case of PSP. Results for synthetic datasets corroborate the trends above (see Supplementary Section 1.4).

### 3.3 MIP Indel solutions are constrained by the indel footprint in extants

Both MIP and BEP find solutions that are based on edges in the POAG, essentially constraining ancestors to exhibit combinations of the same indel patterns that are appear in one or more extant sequences. SICP and PSP performs inference at a different level of resolution; PSP uses patterns of individual gaps present in extant sequences as input, which allow novel indel patterns to occur in ancestors once recast in terms of sequences (which in turn can be recast as edges forming part of a ancestral POAG). Analogously, SICP identifies a set of indel states each of which is inferred independently; their overlap can give rise to novel patterns. Here, we wanted to establish how methods compared in terms of how different ancestors looked relative to extant sequences.

For MIP, the inferred indel edges at an ancestral branch point constitute a subset of the edges present in the input POAG. This constraint ensures that the indel pattern inferred for an ancestor matches the indel footprints observed in the extant sequences. To quantify the degree of a match or (conversely) the distinctiveness of indels at ancestors, we counted ancestral branch points with “new” edges across all datasets (see Figure 5).

**Figure 5:**
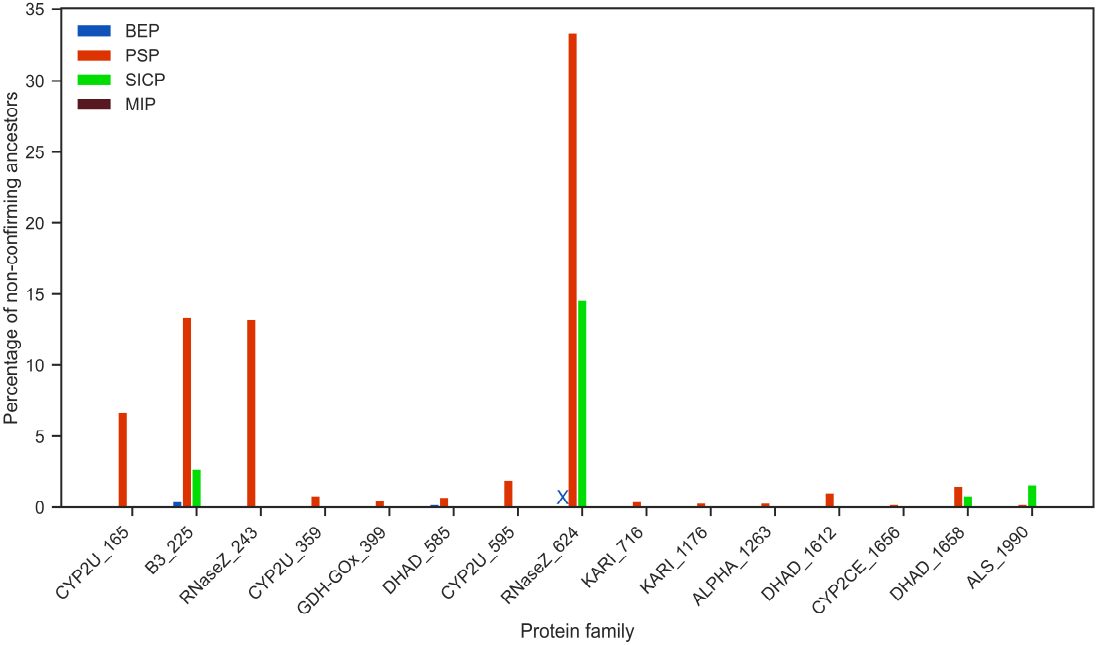
Percentage of non-conforming ancestors for 15 real datasets. X in the above figure denotes that BEP did not provide a solution for the dataset RNaseZ_624.

Unsurprisingly, PSP creates the greatest number of ancestors with novel indel patterns across almost all datasets. SICP is much less creative but forms novel ancestral indels for the most complex datasets.

### 3.4 MIP Indel typically finds alternative optimal indel histories

BEP is able to recover multiple indel histories that explain the same input equally well. The availability of indel options can be used by experimentally resurrect distinct mutants likely to be structurally and functionally viable as demonstrated in [7]. To investigate the hypothesis that phylogenetic trees encompass ambiguous optimal indel histories, we extend the mathematical MIP formulation (Section 2). This extension incorporates an additional constraint (Equation 11), to eliminate the initial optimal solution, allowing the MIP model to identify alternative optimal solutions.

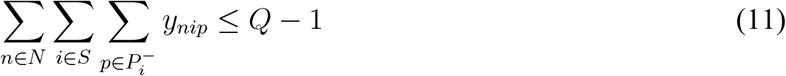

where *Q* is the sum of all the edges present in the first optimal solution.

Out of 15 real datasets, MIP found alternate optimal indel histories for 12 datasets. We investigate the two alternative histories for RNaseZ_243 dataset and examine the indel events in the clade under the ancestral nodes N236 for the RNaseZ_243 tree across both optimal solutions. Figure 6A shows the indel events for the clades under ancestral node N236. While the total number of indel events in the clade under N236 is consistent across both solutions, differences in indel patterns are observed in ancestral nodes N238 and N239 at sites 991 and 992, as illustrated in Figure 6B.

**Figure 6:**
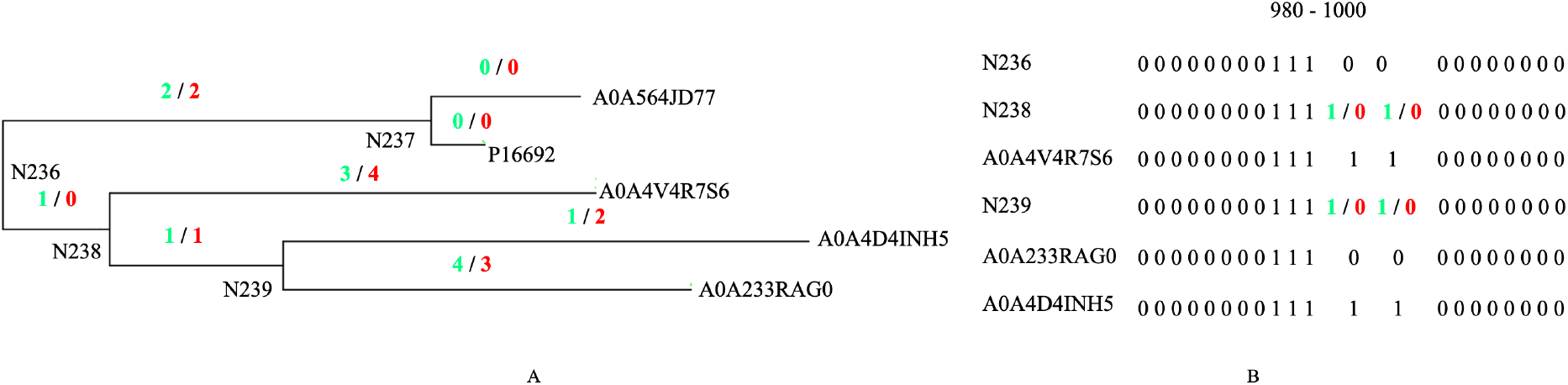
A) The indel events at each branch point for optimal solution one and two (relative to the ancestor) is shown in green and red, respectively. Names starting with N are the ancestral nodes. We note that the total indel events in clade under ancestral nodes N236 is the same in both optimal solutions. The extants are represented by their UniProt identifiers. B) Visualisation of molecular sequence indel patterns for select nodes under the ancestral node N236 for sites from 980 - 1000. Ancestral node N237 is not shown as it has the same indel pattern in both optimal solutions. The sites, 991 and 992, accommodates both gap and ungap scenarios in a parsimonious manner. This is evident by the same sites in the extant sequences A0A233RAG0, A0A4D4INH5 and A0A4V4R7S6 present under the ancestral nodes of N238 and N239.

## 4 Discussion

Indels factor heavily into the adaptive evolution of proteins. Yet, efforts to infer them–some based on evolutionary models [23] and some account for their variable-length sequence imprint [27]–have had limited success in analysing realistically large protein families. The need to discern indel patterns in extant sequences and decode their history is transformed here into a combinatorial but practical optimization problem.

Mixed-Integer programming models are ideal for problems that involve identifying the optimal subset of items (decisions, activities) meeting specific criteria from a well-defined finite set of alternatives. Indel inference in phylogenetic trees, when framed as MIP problem, offers many advantages.

A key benefit of our MIP formulation lies in treating indel inference as a *single, global* optimisation problem. Reassuringly, the MIP approach has the same indel score as other parsimony-based methods for simple and small trees. This is particularly clear in the case of simulated data, where indel histories are well-defined and are not confounded by the sequence alignment step required for real data. Indeed, for simulated data, MIP always finds a solution that is equal or more parsimonious than the true history (which can contain unobservable, composite indel events).

Differences in indel solutions between methods emerge with larger trees, with the MIP approach consistently scoring conservatively lower on indels. MIP preserves the overarching connectedness of the tree, as evidenced by the quality of solutions. We note that comparable methods are led to local forms of optima and show that their combinations sometimes compromise cohesion of indel events when viewed across the whole tree.

The inclusion of constraints in the MIP model plays a crucial role in ensuring plausible histories. For instance, constraints restrict indels at internal branch points from deviating from their neighbours. Inspired by pairwise alignment, the objective function imposes (user-specified) affine gap penalties between ancestors and their direct descendants. We also demonstrate that by formulating constraints around edges in a partial-order graph, the indel events at each ancestor do not deviate from those implied by extant sequences; the edges used to define each ancestor are a simple subset of those that define the input alignment.

The input does not always offer sufficient material to disambiguate solutions; we show MIP can report all indel events that explain the provided evidence. We expect this flexibility can be further harnessed to improve the quality of the underlying MSA, but also help explore diverse sequences and their histories for protein engineering.

Finally, it is worth noting that the MIP approach we present here is the first indel inference method that guarantees a globally optimal solution to realistically large protein family trees; it is applicable to data of the scale that will support a wide range of analyses, e.g. to understand evolutionary drivers of anti-biotic resistance and the engineering of enzymes for biofuel production. Our benchmarking indicates that MIP Indel inference succeeds in finding optimal solutions for trees encompassing over 2000 extant proteins, which is a reasonable time frame for phylogenetic analysis on modest hardware. This efficiency stands in stark contrast to other global optimisation methods like SP, where the time complexity increases exponentially with tree size. For MIP, the number of constraints and variables in the model are proportional to the number of nodes multiplied by the number of sites. Thus, efficiency will deteriorate for large trees; trees with 10,000 sequences are not likely to finish within a few hours without heuristics, which in turn compromises the attractive guarantees of optimality.

## Acknowledgements

We thank Dr Rhys Parry, Anthony Bengochea, and Samuel Davis for providing additional phylogenetic tree protein datasets.

