## Supplementary Material for "Optimal Phylogenetic Reconstruction of Insertion and Deletion Events"

### 1 Supplementary information

#### 1.1 Real datasets

The details of the real datasets used in this paper are shown in Table 1.

| Dataset | Protein family | Total extants | Aligned sequence length |
|---|---|---|---|
| CYP_165 | Cytochrome P450 | 165 | 627 |
| B3_225 | Metallo-$\beta$-lactamase B3 | 225 | 1652 |
| RNaseZ_243 | Metallo-$\beta$-lactamase | 243 | 1213 |
| CYP_359 | Cytochrome P450 | 359 | 542 |
| GDH-GOx_399 | Glutamate dehydrogenase | 399 | 819 |
| DHAD_585 | Dihydroxy-acid dehydratase | 585 | 776 |
| CYP_595 | Cytochrome P450 | 595 | 664 |
| RNaseZ_624 | Metallo-$\beta$-lactamase | 624 | 5240 |
| KARI_716 | Ketol-acid reductoisomerase | 716 | 735 |
| KARI_1176 | Ketol-acid reductoisomerase | 1176 | 513 |
| ALPHA_1263 | Alphavirus | 1263 | 1567 |
| DHAD_1612 | Dihydroxy-acid dehydratase | 1612 | 835 |
| CYP2CE_1656 | Cytochrome P450 | 1656 | 1277 |
| DHAD_1658 | Dihydroxy-acid dehydratase | 1658 | 2525 |
| ALS_1990 | Acetolactate synthase | 1990 | 851 |

Table 1: Real protein families datasets used for MIP Indel evaluation.

The dataset CYP_165, CYP_359, GDH-GOx_399, DHAD_585, CYP_595, KARI_1176, DHAD_1612 are referenced from [7]. The details for the curation of the datasets B3_225, RNaseZ_243, RNaseZ_624, KARI_716, ALPHA_1263, CYP2CE_1656, DHAD_1658 and ALS_1990 is described in the Table 2. In all the datasets, the sequences were aligned using MAFFT [11] and trees were inferred using IQTREE2 [20].

| Dataset | Detail |
|---|---|
| B3_225 | The sequences belong to subclass B3 Metallo-$\beta$-lactamases. We use Glyoxalase II as an outgroup. |
| RNaseZ_243 and RNaseZ_624 | RNaseZ_243 and RNaseZ_624 are Ribonuclease Z sequences across archaea, bacteria, and eukaryotes either limiting the eukaryotic sequences to expected ortholog of the human elaC 1 protein (RNaseZ_243) or also including ortholog of the longer human elaC 2 (RNaseZ_624). We use PhnP sequences, a phosphodiesterase, as an outgroup. |
| KARI_716 | The sequences belong to Class I ketol-acid reductoisomerases. We place the tree root on the branch between the archaeal and bacterial sequences. |
| ALPHA_1263 | The sequences belong to 38 species and 1263 non-redundant sequences from the Alphavirus genus (Family Togaviridae). The tree is rooted at the midpoint. |
| CYP2CE_1656 | The sequences belong to the Cytochrome P450 subfamilies CYP2C and CYP2E. We use vertebrate CYP2 family as an outgroup. |
| DHAD_1658 | DHAD_1658 are dihydroxyacid dehydratases and sugar acid-specific dehydratases from the ilvD/EDD superfamily which includes representation of functionally-distinct clades [16]. We use 6-Phosphogluconate dehydrogenase sequences as an outgroup. |
| ALS_1990 | ALS_1990 are acetolactate synthases and we use a small selection of anabolic acetohydroxy-acid synthases as an outgroup. |

Table 2: Information on the remaining real datasets used in this paper (not referenced from [7]). These datasets are newly created for different protein families.

#### 1.2 TrAVIS synthetic data generation

This section outlines the details around how synthetic data is generated using TrAVIS, which forms part of the GRASP-suite [7]. TrAVIS samples the Gamma distribution $\gamma(\kappa, \theta)$ (where $\kappa \in \{0.5, 1, 2\}$ defines the shape and $\theta = 0.2$ the scale) to set distances on each branch (in turn normalised to the mean $\delta$), bi-furcating each branch point until the specified number of sequences have been mapped as leaves.

For each tree, substitutions, insertions and deletions were randomly introduced at each branch point. Starting at the root with an arbitrary amino acid sequence with a length 80 or longer, the sequence at each of its children is determined recursively. For a sequence, each position is considered as a possible site for a mutation as a Poisson process with a relative rate $2r$ ($r = 1$ for all our synthetic data); following the recipe given by Cartwright [2], $\eta = e^{-2rt}$ is the probability that no indel has occurred, which implies a substitution (determined via a model like that suggested by Le and Gasquel [14], for instance); the probability of an insertion is $(1 - \eta)/2$ which is equivalent to that of a deletion. Finally, the length of the insertion (or deletion) is given by a sampling the Poisson distribution $f(\lambda)$ (where mean $\lambda = 1$, which will give 0 or 1 37% of the time, with greater widths less frequent). Depending on the length of the sequence provided at the root, the resulting aligned sequences ranged from 143 to 2606 positions across the 60 trees.

#### 1.3 Visualisation of indel events

We were curious to understand how methods differ qualitatively, including how indel events distribute across trees. We developed a visualisation where the root node is designated as having zero indels. Following a post-order traversal of the tree structure, we enumerate subsequent events in relation to the root node and cumulatively show their number on the y-axis. The x-axis signifies the cumulative level of the internal node relative to the root at 0. We show the predicted indel events for RNaseZ_243 in Figure 1 illustrating no particular bias and the overall trend that MIP includes least events.

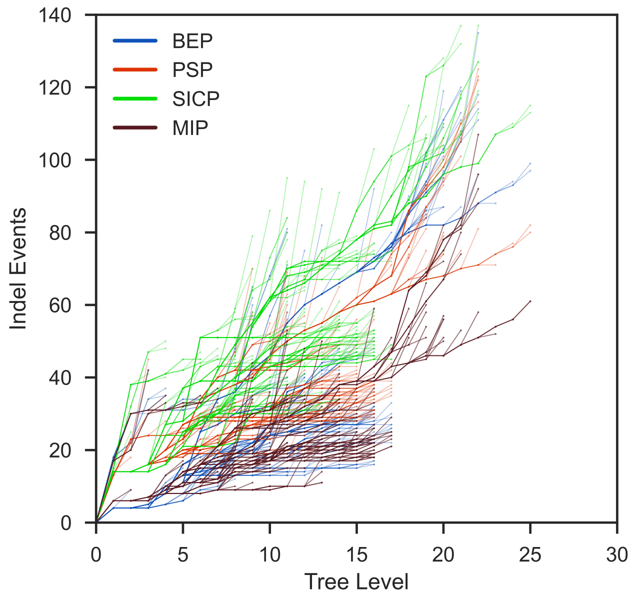


Figure 1: Cumulative indel events predicted (y-axis) for the RNaseZ_243 dataset disperse evenly across different levels of the tree (x-axis) for all methods. MIP tends to predict fewer indels (brown lines).

#### 1.4 Results on synthetic datasets

##### Indel scores for synthetic datasets

Figure 2 shows indel scores for 48 synthetic datasets.

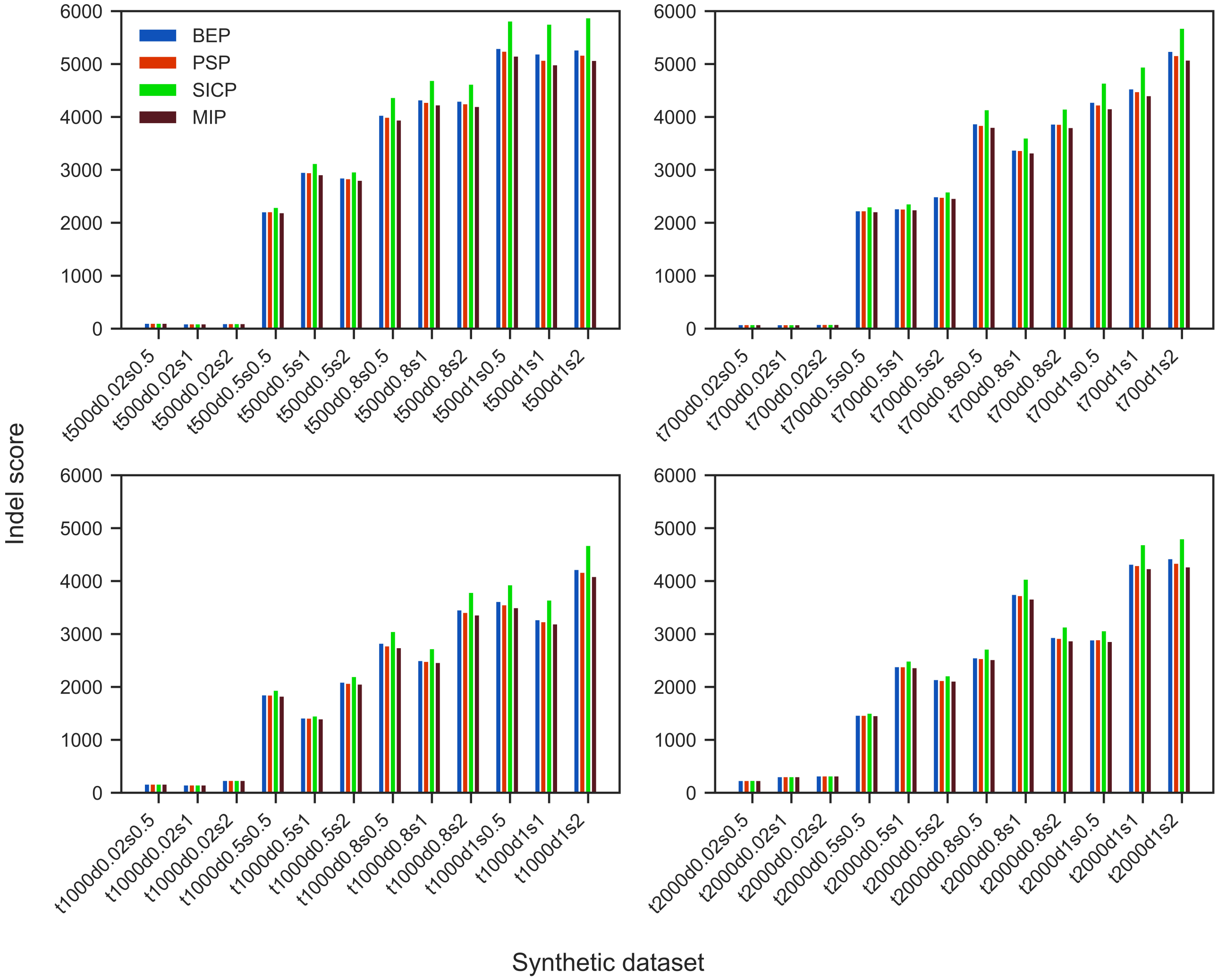


Figure 2: Indel scores for 48 synthetic datasets. Remaining 12 not shown as they report similar trends.

##### Evolutionary cohesiveness in synthetic datasets

Figure 3 shows percentage of branch points where at least one indel state differs from all its descendants and ancestor in 48 synthetic datasets.

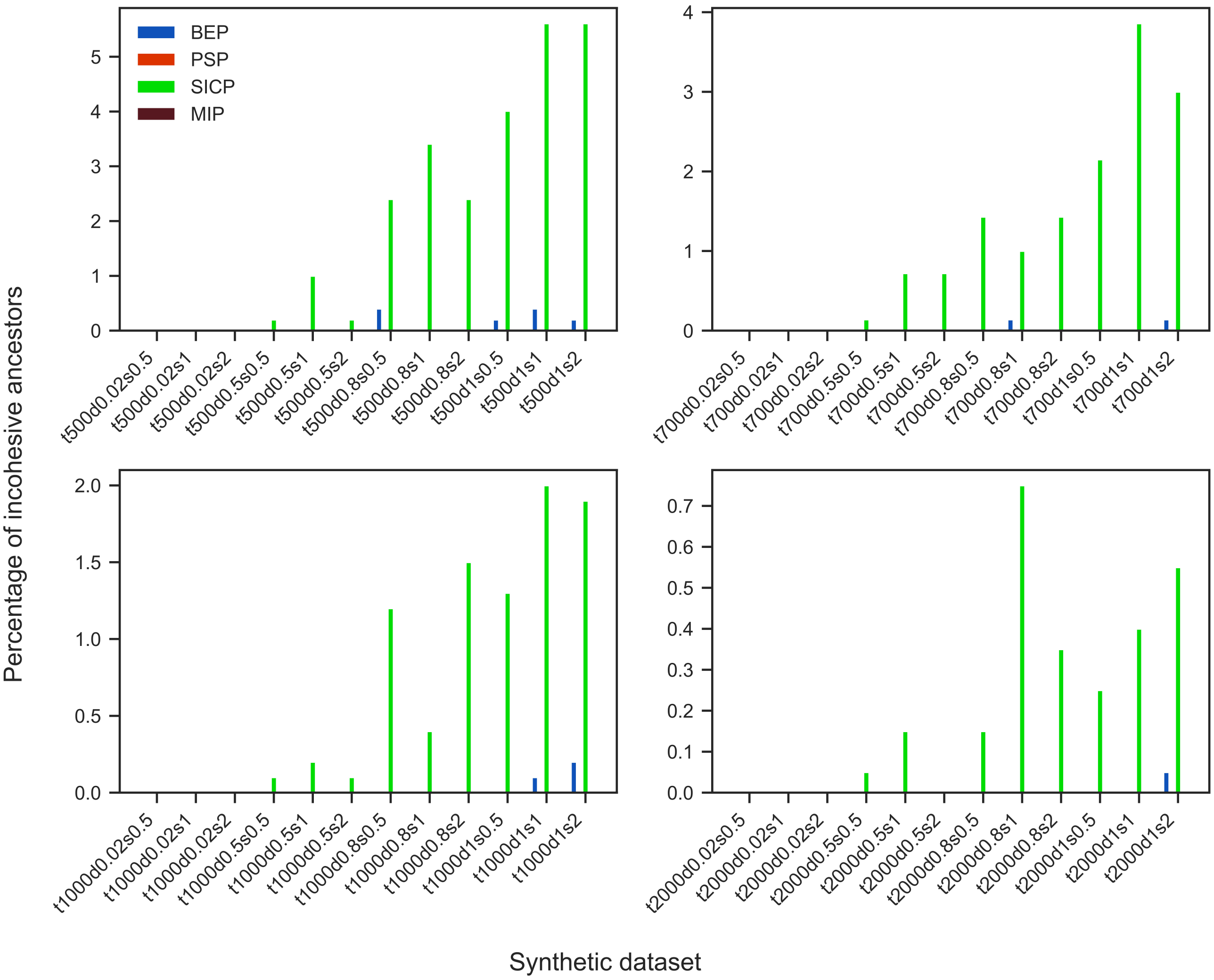


Figure 3: Percentage of incohesive ancestors for 48 synthetic datasets. Remaining 12 not shown as they report similar trends.

##### Indel footprints in synthetic datasets

Figure 4 shows percentage of non-confirming ancestors in 48 synthetic datasets.

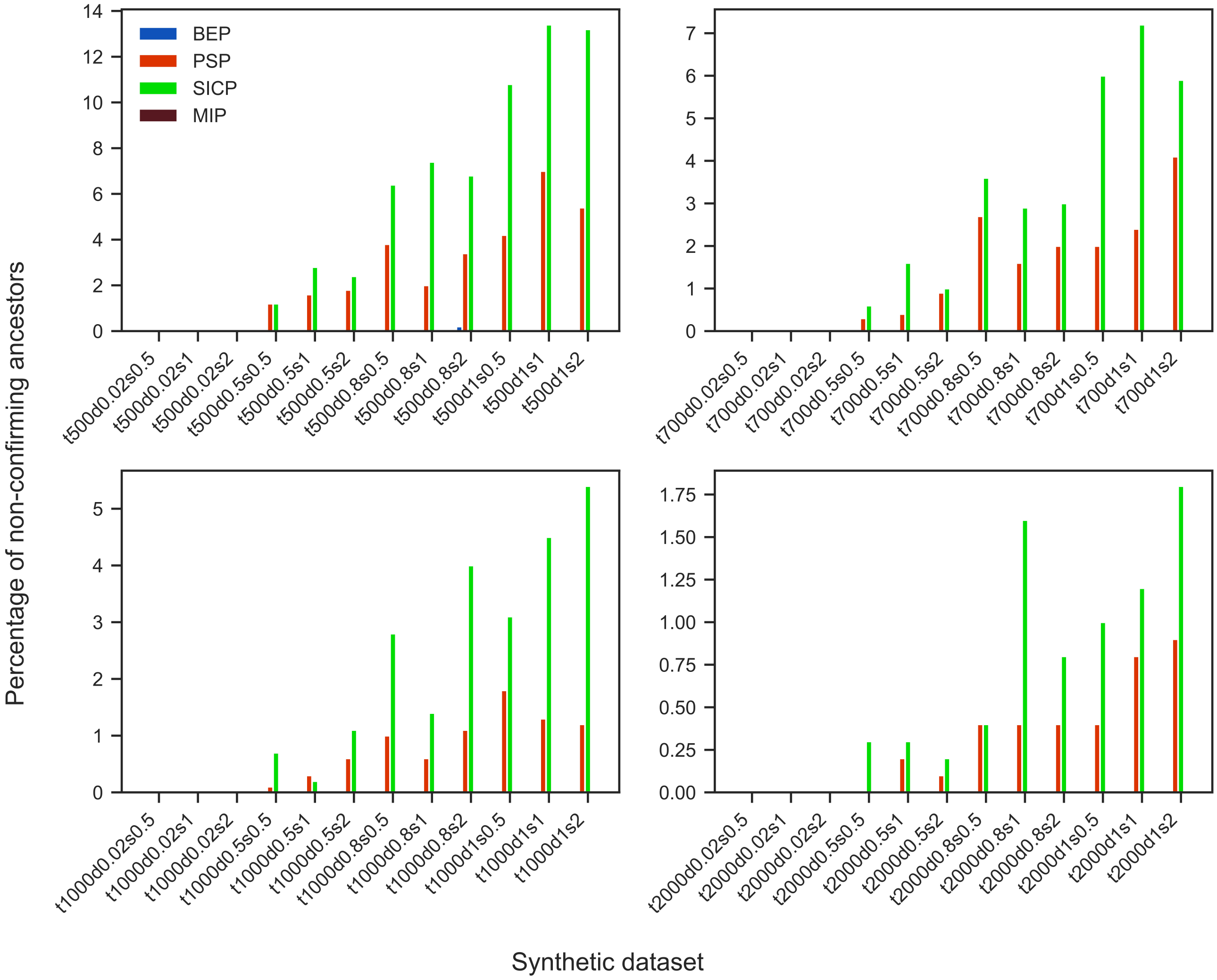


Figure 4: Percentage of non-confirming ancestors for 48 synthetic datasets. Remaining 12 not shown as they report similar trends.

#### 1.5 Hyperparameter experiments

We conducted experiments to gauge the impact of hyperparameter $\alpha$ on indel scores for the synthetic dataset t300d0.5s0.5 (see Figure 5). The indel score peaks when $\alpha$ is zero since there is no penalty for gap opening. As $\alpha$ increases, the indel score decreases, and then levels off.

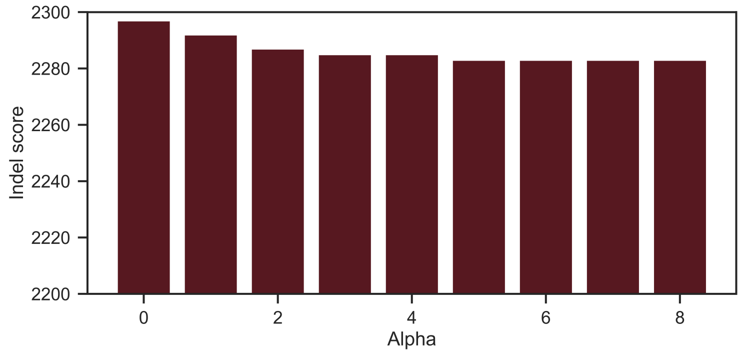


Figure 5: Indel score for different $\alpha$.

#### 1.6 Comparison with Snir and Pachter's method.

Here we present the results of experiments from running indel inference on very small synthetic phylogenetic trees generated by tool TrAVIS using $\delta$ of 1.0 and $\kappa$ of 1.1. We generate datasets by varying the number of extants and sequence positions, assigning them corresponding names. For example, the label `t10p100` denotes a tree with 10 leaves and sequences containing 100 positions. The MIP Indel inference method is compared against the dynamic programming method proposed by [28]. We implemented the SP algorithm and made it available at https://github.com/Tingzhaooo/spindel.

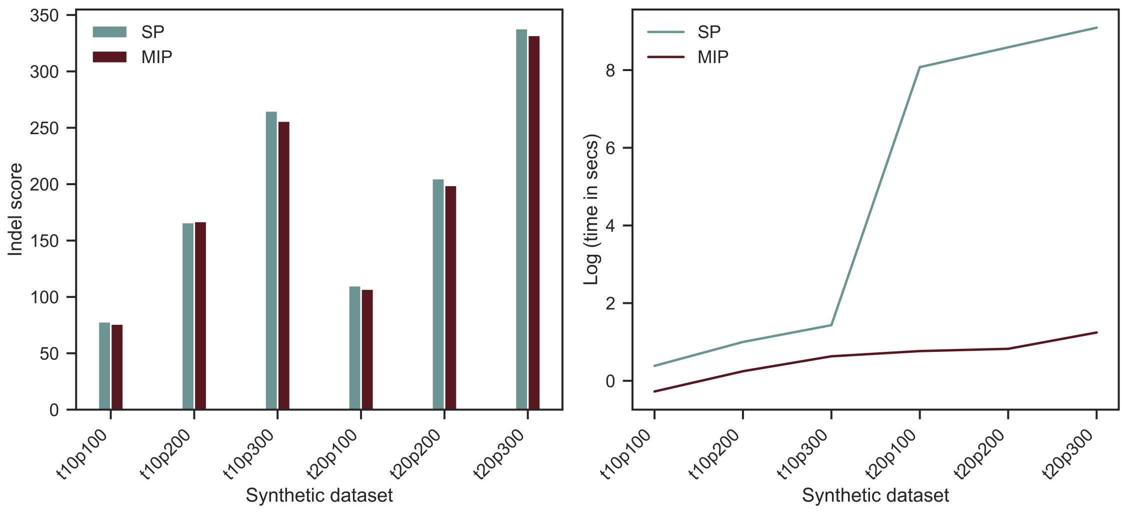


Figure 6: Indel scores and time performance of Snir and Pachter's method (SP) compared with MIP.
